# Thienopyrimidine amide analogs target MmpL3 in *Mycobacterium tuberculosis*

**DOI:** 10.1101/2025.06.26.661674

**Authors:** Vanessa Pietrowski Baldin, Christopher L. Harding, Diana Quach, Joseph Sugie, Joe Pogliano, Tanya Parish

## Abstract

**Objectives:** The identification of novel agents with mechanisms of action distinct from those currently utilized in tuberculosis treatment remains a significant challenge. The mycobacterial protein MmpL3 has emerged as a promising drug target due to its essential role in the synthesis of the cell wall of *Mycobacterium tuberculosis*. We previously identified novel thienopyrimidine amides with good anti-tubercular activity.

**Methods:** We profiled a subset of thienopyrimidine amides determining activity against intracellular bacteria and bactericidal activity against replicating bacteria. We ran assays to determine mode of action by measuring cell wall stress, ATP production, and bacterial cytological profiling. We determined activity against a strain of M. tuberculosis with mutations in MmpL3. We isolated and sequenced resistant mutants.

**Results:** We tested five analogs against a strain of *M. tuberculosis* with mutations in MmpL3 and determined that they lost potency. Analogs induced P_iniBAC_, a reporter for cell wall stress, and led to an ATP boost characteristic of cell wall inhibitors. Bacterial cytological profiling of a representative compound revealed a morphological profile consistent with other MmpL3 inhibitors.

**Conclusions:** Together, our data support MmpL3 as the most probable drug target for the TPA analogs and add to the growing list of scaffolds that can inhibit this vulnerable transporter.

## Introduction

Despite the availability of drug treatment regimens, tuberculosis remains a major global health concern, with an estimated 8.2 million new cases reported in 2023, including 400,000 cases of drug resistant tuberculosis (1). Currently, no vaccine provides effective protection against pulmonary TB. Additionally, the bacteria can enter a latent phase, where they display no symptoms, or cause symptoms that mimic other pathologies (2). Together, these factors highlight the urgent need for intensified efforts in TB control and the development of novel therapies, particularly those targeting drug-resistant bacilli and aiming to shorten the duration of treatment.

*Mycobacterium tuberculosis* is the causative agent of TB in humans. *M. tuberculosis* has a thick cell wall, which is composed by a highly hydrophobic bilayer of mycolic acids linked to arabinogalactan in turn linked to peptidoglycan. This structure forms a unique barrier against drugs and the host immune defenses (3–5). Hence, cell wall biosynthesis represents an attractive target for the development of new inhibitors.

MmpL3 (Rv0206c) is an essential transporter which is highly conserved across the *Mycobacterium* genus (4, 6, 7) and is the only member of the MmpL family that is essential in *M. tuberculosis* (2, 4, 7). Its essentiality is attributed to its role in the translocation of trehalose monomycolate (TMM) to the periplasmic space, where it serves as a precursor for the synthesis of trehalose dimycolate (TDM), a critical component of the cell envelope (4, 8). Given the important role of MmpL3 in constructing the mycobacterial cell wall, it has emerged as a high-value, druggable target, and at the moment, one of the most studied for anti-TB drug development (6). There are numerous MmpL3 inhibitors with diverse chemical structures which have potent antibacterial activity (5, 9).

We previously identified and characterized novel thienopyrimidine amide (TPA) analogs that inhibit *M. tuberculosis* growth and had low cytotoxicity. From this series, we identified two subsets. One subset (TPA-L) had increased activity against a LepB (signal peptidase) hypomorph, indicating that the signal peptidase or protein secretion is the target or mode of action (10, 11). The second subset of molecules (TPA-M) had excellent potency but were equipotent against wild-type and hypomorph strains of *M. tuberculosis* suggesting that the target was not protein secretion. In this paper we explored the biological profile of the TPA-M series of molecules and identified MmpL3 as the most likely intracellular target.

## Material and Methods

### Bacterial strains and culture conditions

*M. tuberculosis* H37Rv-LP ATCC 25618 (wild-type) (12) and *M. tuberculosis* LP-0497754-RM301 (MmpL3 F255L, V646M, F644I) (13) were cultured in Middlebrook 7H9 broth medium supplemented with 10% oleic acid, albumin, dextrose, and catalase (OADC) enrichment (BBL/Beckton and Dickson) and 0.05% w/v Tween 80 (7H9-Tw-OADC). Nutrient-starved cells were generated by resuspending cells in phosphate-buffered saline (PBS) with 0.05% w/v Tyloxapol (PBS-Tyl) at OD 1.0 and incubating for 7 days at 37°C without agitation. Avirulent *M. tuberculosis* mc^2^6206 (Δ*pan*CD Δ*leu*CD) was grown in 7H9-Tw-OADC supplemented with 0.5% wt/vol glycerol, 0.2% wt/vol Casamino Acids, 48 µg/mL pantothenate, and 50 µg/mL leucine at 30°C.

### Determination of minimum inhibitory concentration (MIC)

TPA molecules (**Figure 1**) were synthesized as previously described (10). Compound stocks were resuspended in DMSO and stored at -20°C. MICs were determined in 96-or 384-well microplates as described (14). Briefly, *M. tuberculosis* was grown to mid-log phase; cultures were dispensed into plates containing test compounds to a final OD_590_ of 0.02 and incubated at 37°C for 5 days (H37Rv) or 7 days (mc^2^6206). The OD_590_ was measured and IC_90_ determined as the concentration at which 90% of growth was inhibited as compared to controls (DMSO only).

**Figure 1.**
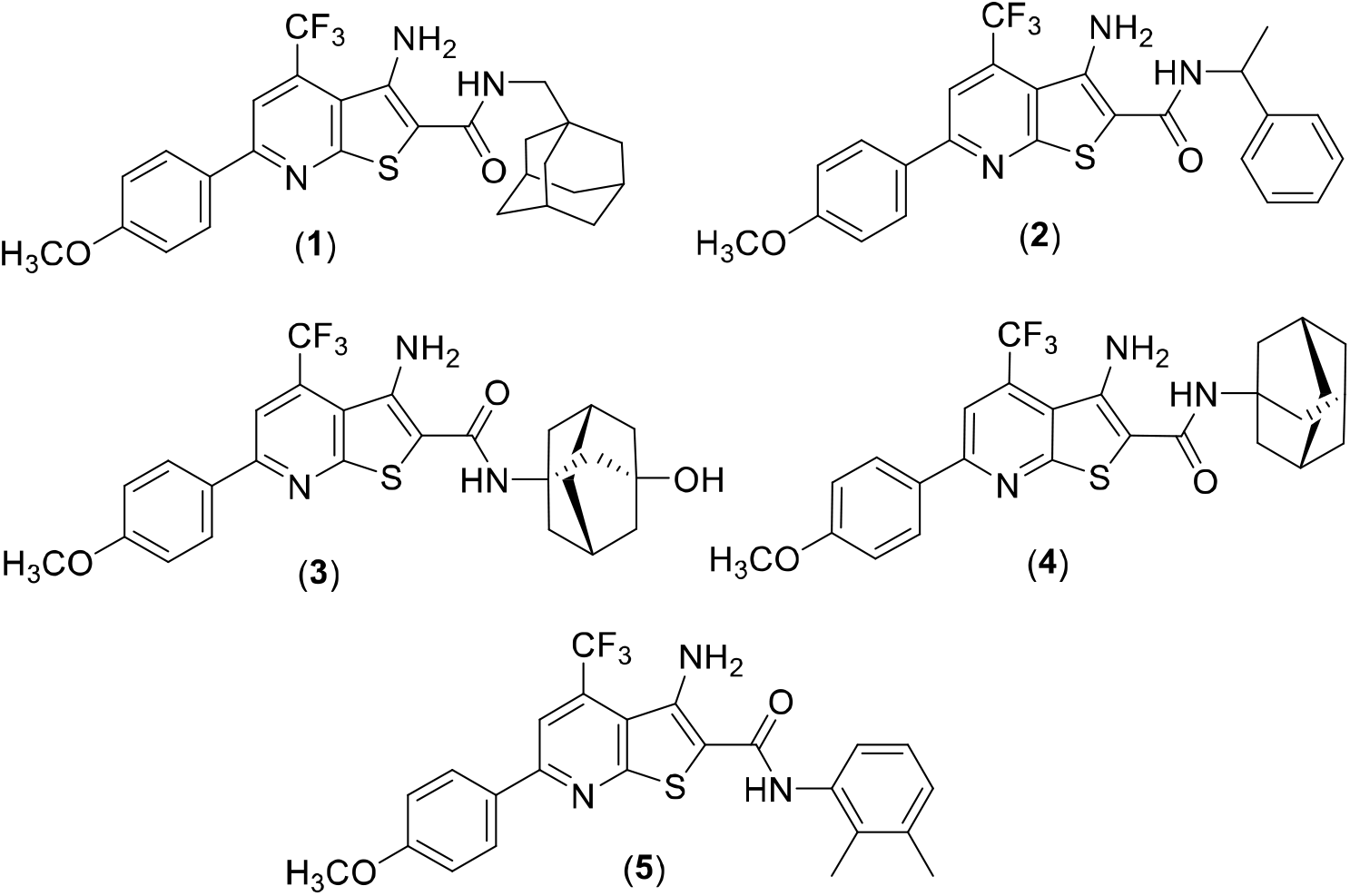
Structure of analogs used in this study. (1) TPN-0102024 (2) TPN-0099994 (3) TPN-0099934 (4) TPN-0099730 (5) TPN-0089300

### Determination of activity against intracellular bacilli

THP-1 cells were cultivated in RPMI-1640 medium supplemented with 10% FBS and incubated at 37°C, 5% CO_2_. Cells were treated with 80 nM PMA for 24 h prior to infection, harvested using Accumax™ solution, and resuspended in fresh cRPMI at a final density of 9×10□ cells/mL. Cells were infected overnight at a multiplicity of 1:1 with *M. tuberculosis* expressing LuxABCDE and exposed to compounds in 96-well plates for 72 h at 37°C and 5% CO_2_. Bacterial viability was assessed by luminescence; IC_50_ was determined as the concentration at which 50% of growth was inhibited as compared to controls (DMSO only).

### Determination of bactericidal activity

*M. tuberculosis* H37Rv-LP was cultured in 7H9-Tw-OADC or starved for 7 days in PBS-Tyl, adjusted to a theoretical OD_590_ of 0.02 and exposed to compounds in 96-well plates. Cultures were spotted onto 7H10 agar supplemented with v/v 10% OADC (7H10-OADC) on day 0, 7 and 14. The MBC was defined as the lowest concentration at which no visible growth was observed. Niclosamide was included as a control.

### Induction of cell wall stress

*M. tuberculosis* strain carrying the P_iniBAC_-lux reporter plasmid (15) was grown to mid-log in GAST/Fe protein-free medium containing 0.3 g/L Bacto Casitone, 0.05 g/L ferric ammonium citrate, 4 g/L dibasic anhydrous potassium phosphate, 2 g/L citric acid, 1 g/L L-alanine, 1.2 g/L magnesium chloride, 0.6 g/L potassium sulfate, 2 g/L ammonium chloride, 1.8 mL of 10 M sodium hydroxide, 10 mL glycerol, 5 mL 10% Tween 80, plus 15 µg/mL kanamycin. Cultures were adjusted to a theoretical OD_590_ of 0.02 and used to inoculate 96-well plates containing compounds. After 72 hours of incubation at 37°C, 1 vol of 10 mg/mL luciferin in of 1M HEPES buffer pH 7.8, 1M DTT was added and RLU was read after 25 min of incubation at RT. Ethambutol was included as a positive control.

### Determination of ATP levels

ATP was measured using the BacTiter-Glo Assay kit (Promega) according to the manufacturer’s instructions. Log phase *M. tuberculosis* was exposed to compounds for 24 h, .50 µL BacTiter-Glo™ was added, incubated at RT for 10 min and RLU read. Growth was measured after 120 h by OD. Q203, was included as a positive control.

### Isolation of resistant mutants

Log phase *M. tuberculosis* was plated on 7H10-OADC plates with 5X solid MIC and 10X solid MIC. Plates were incubated at 37°C until isolated colonies appeared. Potential resistant mutants were picked and streaked onto 5X MIC. MIC in liquid medium was measured to confirm resistance. Genomic DNA from three isolates was extracted from cultured cells by heat inactivation for 10 min at 100°C followed by 0.22 µm filtration. PCR amplification was performed using 10 µL of extract DNA in a final volume of 50 µL containing 1 µL of Pfu polymerase (Agilent), 5 µL of 10X Pfu amplification buffer (Agilent), 2.5 µL of primers MmpL3_seq_1_fwd (5’-gattcgctacctgagcag-3’) and MmpL3_seq_11_rev (0.5 µM) (5’-catttactgcagccgctg-3’) and 4 µL of 10 mM dNTP mix. PCR amplification was conducted, products were purified using the Qiagen PCR purification kit, and sequencing was performed by Plasmidsaurus using Oxford Nanopore Technology with custom analysis and annotation.

### Bacterial Cytological Profiling (BCP)

*M. tuberculosis* mc^2^6206 for bacterial cytological profiling was prepared as described (16). Briefly, cultures were adjusted to an OD_600_ of ∼0.06-0.08 and incubated at 30°C for 18-20 hours before exposure to compounds at 1X and 5X MIC for 48h and 120 hours. Cells were fixed using a mixture of 100 µL of 16% paraformaldehyde, 3 µL of 8% glutaraldehyde, and 20 µL of 0.4 M phosphate buffer pH 7.5, washed twice with 200 µL of warm medium, concentrated to approximately 30 µL, and stained for 30 min. Full-field fluorescence microscopy images of the samples were preprocessed from their original proprietary microscope file format into a common image format (TIFF). The original image dimensions (3×2048×2048) were cropped by trimming 124 pixels from each edge to avoid optical artifacts, resulting in dimensions of 3×1800×1800. A 600-pixel square sliding window was passed over this image with a step size of 60 pixels to produce 400 sub-images, which were each passed to a trained convolutional neural network (CNN). Each full-field image, consisting of 400 sub-images, generated a vector that describes a point in latent space. Similarity scores were calculated by comparing the location of unknown compound-treated cells in this space to control compound-treated cells using a scaled version of the average minimum distance.

## Results and Discussion

### TPA analogs are active against intracellular mycobacteria

We previously demonstrated that the TPA series of compounds are active against aerobically-grown *M. tuberculosis* (10). A subset of analogs had potent anti-tubercular activity. Since these analogs had equipotent activity against wild-type and LepB hypomorph strains, we hypothesized that they did not target protein secretion. We wanted to determine the mode of action and target of these potent molecules.

We had already noted that compounds were active against replicating bacteria, so we expanded our work to look at activity against intracellular bacteria and non-replicating bacteria. We selected several active analogs (Figure 1). Molecules were tested for activity against intracellular *M. tuberculosis* in THP-1 cells. All compounds were active against intracellular *M. tuberculosis*, with activity similar to the MICs (Table 1).

**Table 1.**
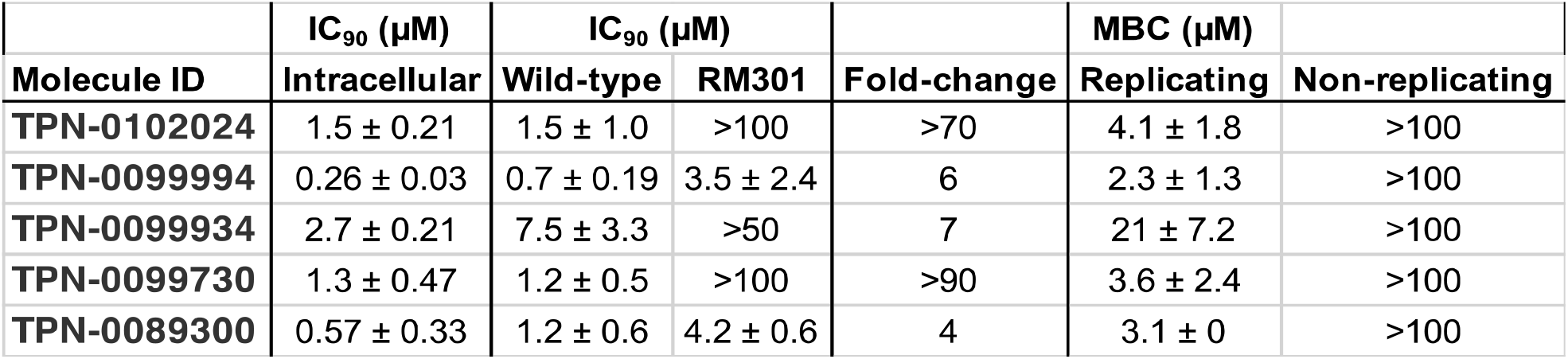
Activity of TPA analogs against *M. tuberculosis*. Intracellular IC_90_ = concentration that leads to 90% inhibition of bacterial replication in THP-1 macrophages. Data are the average and standard deviation of two independent replicates. IC_90_ = concentration that leads to 90% inhibition of bacterial growth in aerobic culture. Data are the average and standard deviation of a minimum of 3 replicates. Fold-change = comparison of IC_90_ for wild-type and RM301 (MmpL3 F255L, V646M, F644I) strains. MBC = minimum bactericidal concentration – concentration at which no viable bacteria were detected after 14 days. Data are the average and standard deviation of three replicates.

### TPA analogs are bactericidal against replicating bacilli

We assessed the bactericidal activity of the TPA analogs against *M. tuberculosis* wild type following a 14-day exposure under replicating conditions as well as against non-replicating bacteria generated by nutrient starvation. All analogs were bactericidal against replicating *M. tuberculosis* with MBC/MIC ratio of <4. No bactericidal activity was seen against non-replicating cells (MBC > 100 µM) (**Table 1**). These data are in contrast to what we previously noted with inhibitors of LepB which showed bactericidal activity against non-replicating bacteria further supporting a different target or mode of action (17).

### TPA analogs induce cell wall stress in *M. tuberculosis*

We were interested in determining the mode of action of our molecules. As a first step, we determined whether molecules induced cell wall stress. We used a reporter strain of M. tuberculosis expressing luciferase under the control of P_iniBAC_, since Induction of *iniBAC* expression is a marker of cell envelope stress (18). We saw induction of PiniBAC for all analogs, which predominantly occurred at concentrations close to the MIC. These data suggest that TPA analogs cause cell wall stress (Figure 2 and Figure S1).

**Figure 2.**
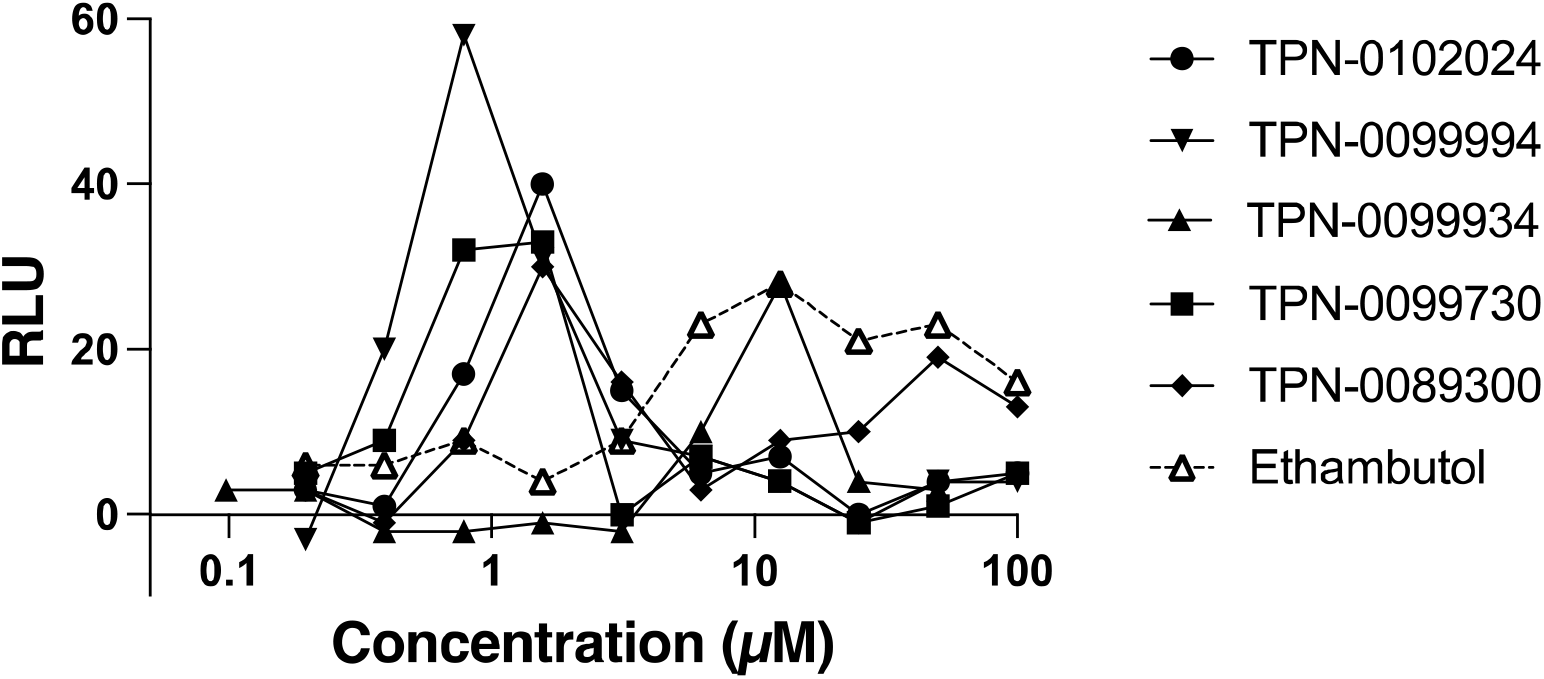
Exposure to TPA analogs induces cell wall stress in *M. tuberculosis*. *M. tuberculosis* P_iniBAC-_Lux was exposed to compounds for 72h and luminescence was read. Data are representative of two independent experiments (see Fig S1).

### TPA analogs boost ATP in M. tuberculosis

A link between induction of *iniBAC* and a burst in ATP production has previously been proposed (19). To determine if our molecules also affect ATP levels, we measured intracellular ATP levels following treatment with TPA analogs. As anticipated, increasing compound concentrations led to a boost in ATP levels (Figure 3 and Figure S2). Again we saw the boost occurring at concentrations around the MIC. These data are consistent with the response seen with other MmpL3 inhibitors (3, 20).

**Figure 3.**
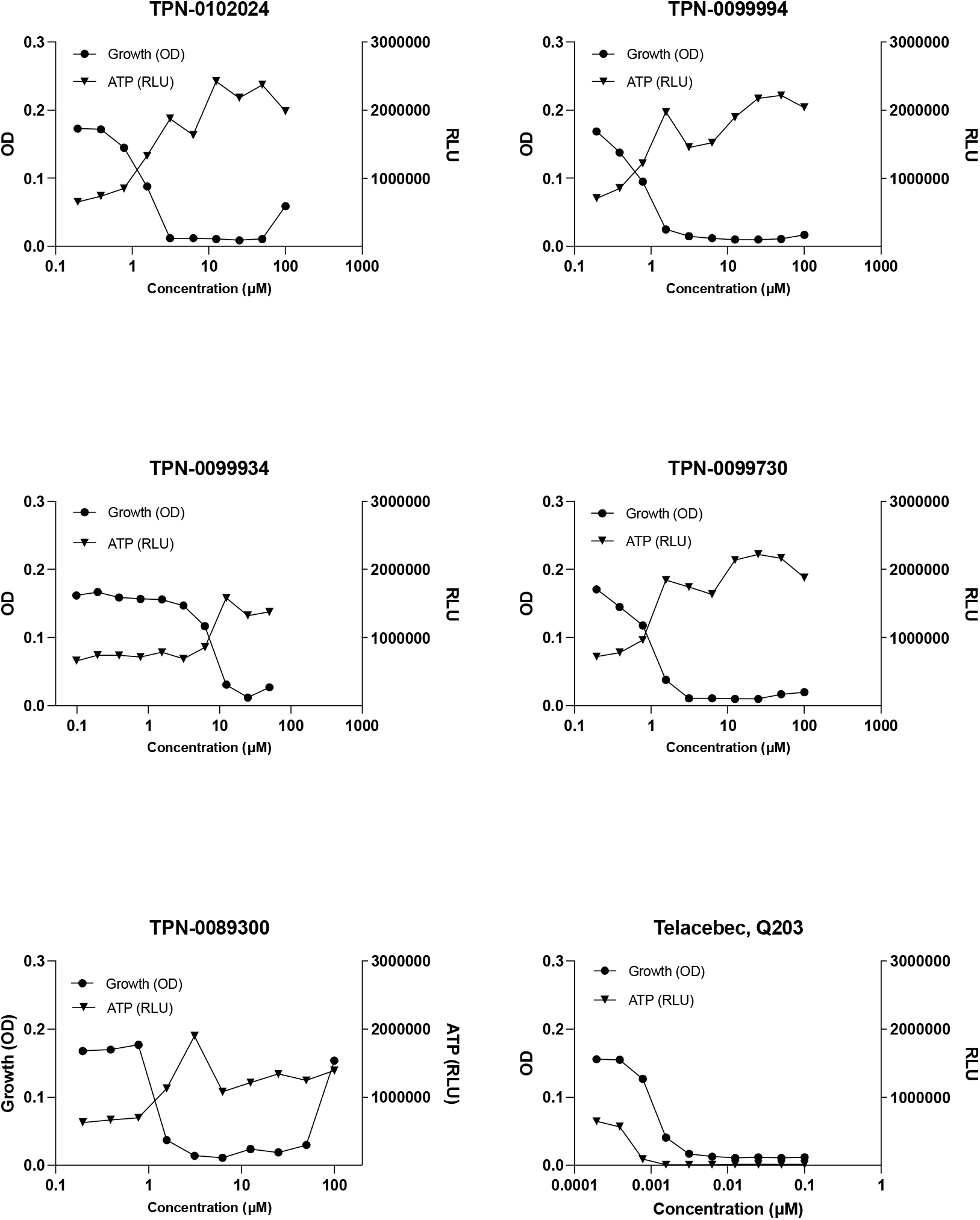
TPA analogs boost ATP in *M. tuberculosis*. *M. tuberculosis* was exposed to compounds for 24h and ATP was measured using BacTiter-Glo. Growth was measured by OD after 5 d. Data are representative of two independent experiments (see Fig S2). Q203 was used as a control.

### MmpL3 mutation leads to resistance to TPA analogs

We hypothesized that our molecules could be targeting MmpL3, since this is a highly promiscuous target and cell wall stress and elevated intracellular ATP levels are features associated with MmpL3 inhibition in *M. tuberculosis* (3). We compared TPA activity against the wild-type *M. tuberculosis* and strain with mutations in MmpL3 which is resistant to a wide range of MmpL3 inhibitors (13, 20). The MmpL3 mutant strain demonstrated increased resistance to all analogs, with at least a 4-fold increase in IC_90_ (Table 1). Interestingly, this strain was highly resistant to three of the five analogs which lost all activity (IC_90_ >50-100 μM). Thus our data are consistent with MmpL3 being the drug target for this compound series.

### Resistant strains have a non-synonymous mutation in *mmpL3*

In order to determine whether there might be additional targets, we isolated resistant mutants using TPN-0089300. Three strains were selected with an IC_90_ of >100 μM. Since MmpL3 mutations lead to resistance, we sequenced *mmpL3* in three resistant isolates (RM3, RM4, and RM6). All isolates had the same non-synonymous mutation in MmpL3 of F644L. This mutation has been associated with resistance to other MmpL3 inhibitors and is similar to one of the mutations in the MmpL3 mutant strain we used to test for resistance (F644I) (4, 8, 13, 21).

### Cytological profiling

We determined the phenotypic effect of a representative TPA analog using bacterial cytological profiling (16). We confirmed that TPN-0089300 was active against the avirulent *M. tuberculosis* strain mc^2^6206 (MIC of 1 µg/mL). Bacterial cells were exposed to TPN-0089300 for 48 and 120 hours at 2X and 5X MIC (2 µg/mL and 5 µg/mL respectively). Morphological profiling demonstrated that bacterial morphology changed with cells becoming more rounded and losing membrane integrity after 120h (Figure 4). Based on the similarity score, this profile was identified as a strong match to other MmpL3 inhibitors (Figure 5) (16, 20).

**Figure 4.**
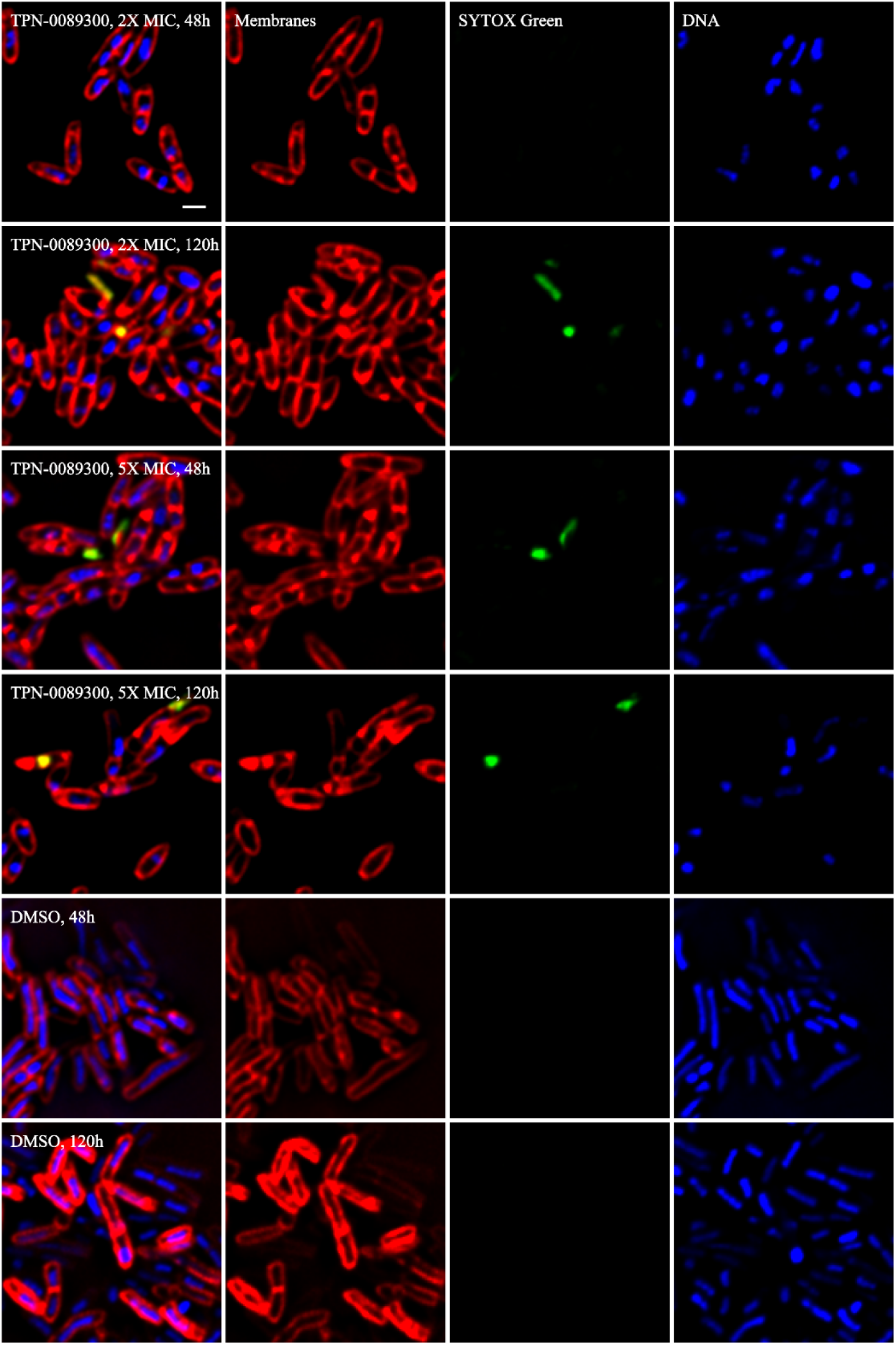
Cytological profiling in response to TPN-0089300. *M. tuberculosis* mc^2^6206 was treated for 48 and 120 h with TPN-0089300 or DMSO (control) and stained. Membranes are shown in red, DNA in blue and green indicates loss of membrane integrity.

**Figure 5.**
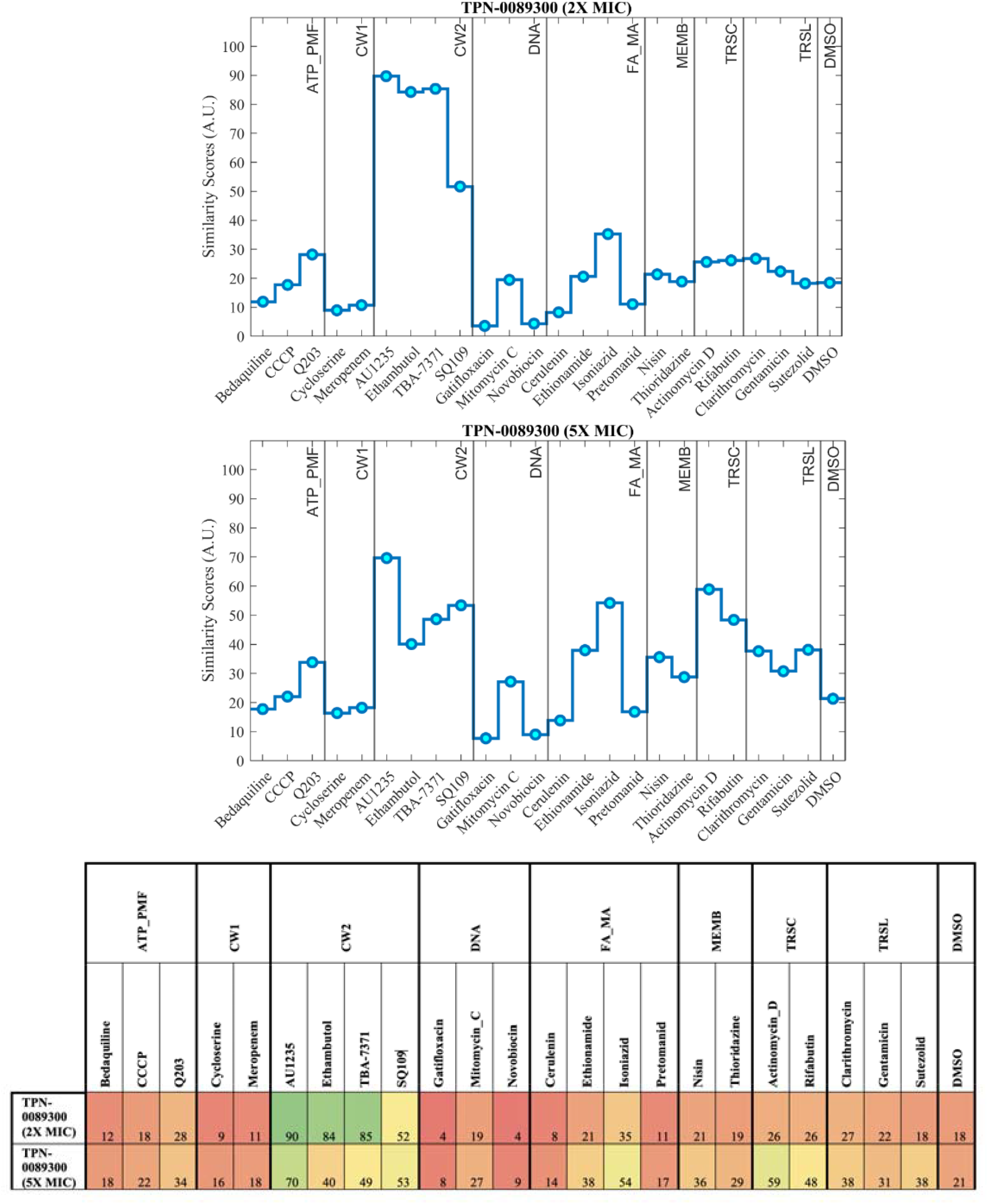
Similarity scores for cytological profiling in response to TPN-0089300.

## Conclusions

We provide strong evidence that a subset of TPA analogs, which we now term TPA-M, target MmpL3 in *M. tuberculosis*. TPA-M compounds were bactericidal for replicating bacilli, induced cell walls stress and boosted ATP production. Mutations in MmpL3 conferred resistance and bacterial cytological profiling identified MmpL3 as the most likely mode of action. We demonstrated that the TPA-M analogs do not kill starved, non-replicating bacilli. This is consistent with other MmpL3 inhibitors that have little effect against non-replicating *M. tuberculosis* (3, 22). Cells treated with MmpL3 inhibitors develop dimples at the dividing pole, where the protein localizes during active cell division and the lack of activity against non-replicating cells has been attributed to the absence of MmpL3 localization at the pole in the non-replicating state (7, 23). Taken together our data suggest that MmpL3 is the primary target of the TPA-M series.

## Acknowledgments

The authors kindly thank Grace Liu, Renee Allen, and Lauren Ames for technical assistance.

