## Supplementary data for "Thienopyrimidine amide analogs target MmpL3 in *Mycobacterium tuberculosis*"

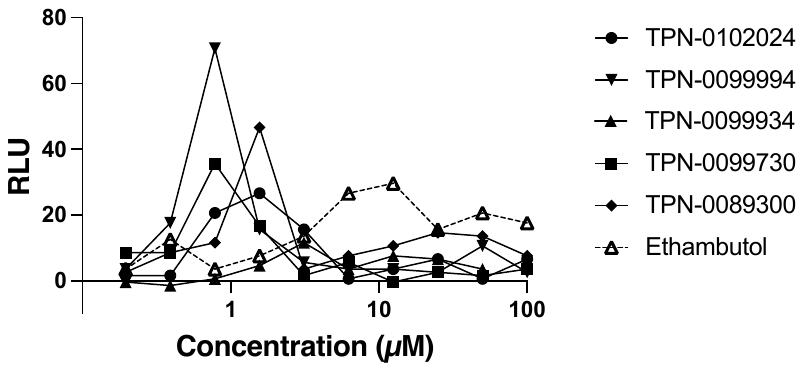

**Figure S1.** **Exposure to TPA analogs induces cell wall stress in *M. tuberculosis*.**

*M. tuberculosis* P_iniBAC-_Lux was exposed to compounds for 72h and luminescence was read.

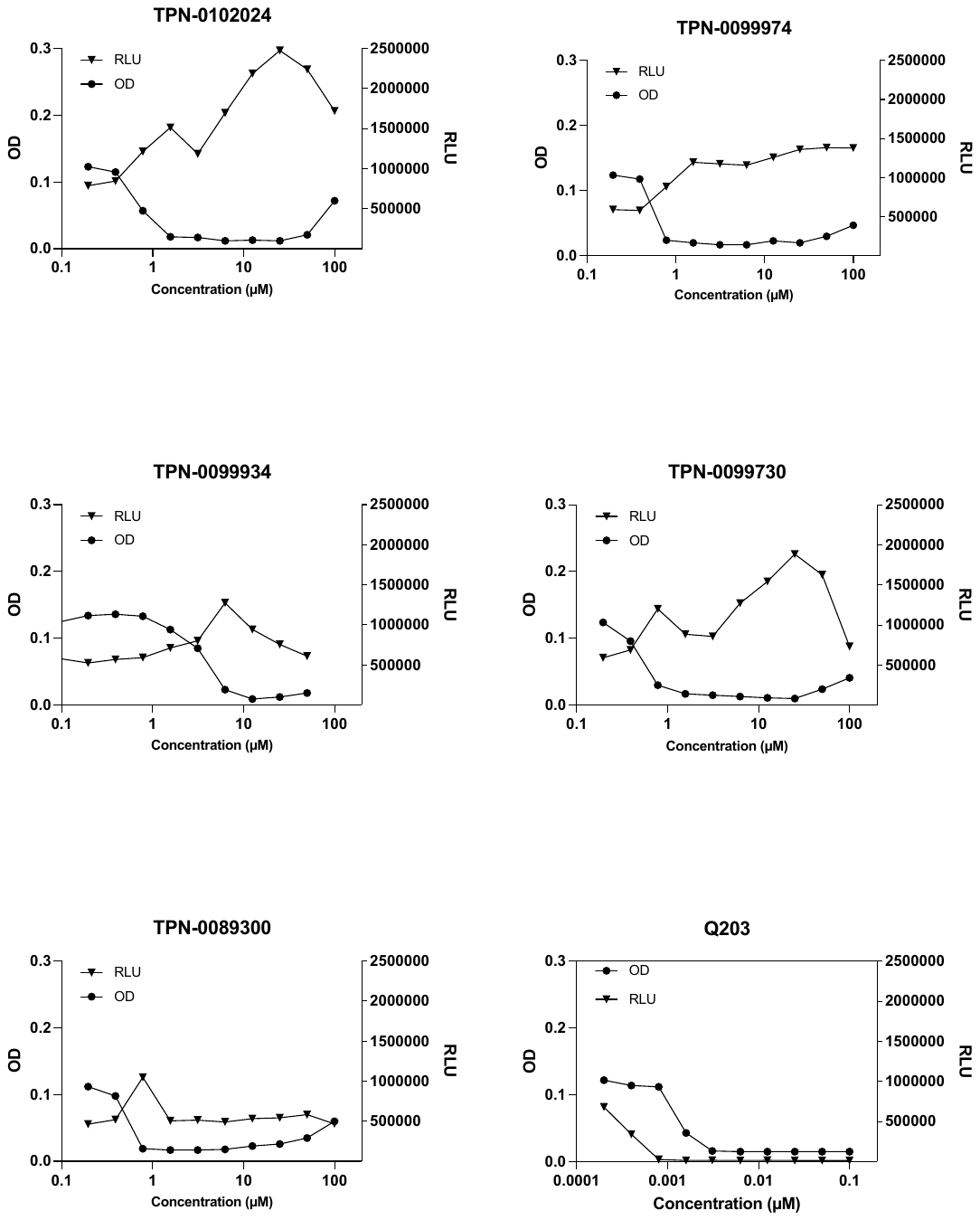

**Figure S2. TPA analogs boost ATP in *M. tuberculosis*.**

*M. tuberculosis* was exposed to compounds for 24h and ATP was measured using BacTiter-Glo. Growth was measured by OD after 5 d.
